# Newly obtained genome of fungi-related amoeba is enriched with genes shared with animals-related protists

**DOI:** 10.1101/2023.12.07.570557

**Authors:** Igor R. Pozdnyakov, Evgeny V. Potapenko, Vera A. Kalashnikova, Christina O. Barzasekova, Vasily V. Zlatogursky, Alexey O. Selyuk, Ksenia M. Sukhanova, Vladislav V. Babenko, Daria I. Boldyreva

## Abstract

Nuclearariids are a group of Opisthokonta, forming the deepest branch in Holomycota - one of the two major Opisthokonta clades, containing Fungi as a crawn group. They are the only members of Holomycota retaining the filose amoeboid state ancestral for Opisthokonta. The newly assembled genome of *Nuclearia thermophila* (Holomycota, Rotosphaerida) had a total length of 49 Mb, 15 321 protein-coding genes and a GC percentage of 44%. This is the first sequenced genome for this genus and the the third for Rotosphaerida as a whole. It was shown that N. thermophila shares more protein domains with Holozoa, than with the rest of Holomycota. Protein domains that were presumably acquired or lost by the common ancestors of the Holomycota and Holozoa groups were identified. The Holomycota ancestor had probably more gains and losses of protein domains compared to the Holozoa ancestor, which is particularly true for metabolism-related domains. However, this trend should be confirmed by studying the genomes of free-living organisms of the Teretosporea group.

## Introduction

Nucleariids, known since the mid-19th century. (Tsenkovsky, 1865), are free-living filose amoebae and represent the deepest branch of fungi-containing Holomycota – the sister group of animals-containing Holozoa (Fig. 1) (Brown et al., 2009; Liu et al., 2009 Gabaldo et al., 2022).. Unlike other more derived members of holomycota, nucleariids uniquely retained free-living predatory lifestyle, while the rest of this clade representatives swithched to being parasites, parsitoids or osmotrophic feeders (Tedersoo et al. 2019). This means that they might hold the keys to understanding the ancestral state of Holomycota and the subsequent modifications, leading to the emergence of fungal kingdom. At the same time this clade lost the ability of eukaryotic flagellum assembly and turned to filose amoeboid condition throughout the whole life cycle (Gabaldón et al., 2022). Remarkably the similar transition to solely filose amoeboid state took place pararelly in filasterians, belonging to Holozoa – the sister lineage of Holomycota (Ros-Rocher et al., 2021). The independedntly acquired similarity is so drastic, that initially *Capsaspora owczarzaki* (the model filasterian) was mistakenly described as nucleariid (Hertel et al., 2002). The transition to solely amoeboid condition and loss of flagellum definitely was accompained with some amount of reductive evolution, which would be interesting to track on the genome level. All of the above makes nucleariids the promising model for reconstructing the ancestral traits of Opistokontha, which were the basis for evolution of major kingdoms of animals and fungi, as well as for studying the reductive process taking place during evolutionary transition to amoeboid state. Despite their obvious value, nucleariids remain poorly studied genetically. Only in recent years, few genomic, metagenomic, and transcriptomic studies have been carried out (Cantor 2015, Galindo and Torruella 2019, Ocana-Pallares et al. 2022). However, the data obtained are mostly present and available in a form of unannotated metagenomes (Torruella 2017, 2019). Successful genome assembly was performed only for Fonticula alba (NSBI accession number GCA_000388065.2), likely a highly derived and secondarily modified slime mould. Apart from this, there is a genomic assembly of Parvularia atlantis (NSBI accession number GCA_943704415.1), which is characterized by low connectivity. Here we present the newly assembled high quality genome of *Nuclearia thermophylla*, a typical free-living bacterivorous representative of nucleariids. Based on the assembled and annotated genome, we carried out a comparative analysis of the protein domains composition. The performed comparative analysis sheds some light on physiological and cytological differences in the early evolution of both Holomycota and Holozoa and reveals the main traits, which shaped the genomic landscape of these two majors Opisthokonta lineages.

**Figure 1.**
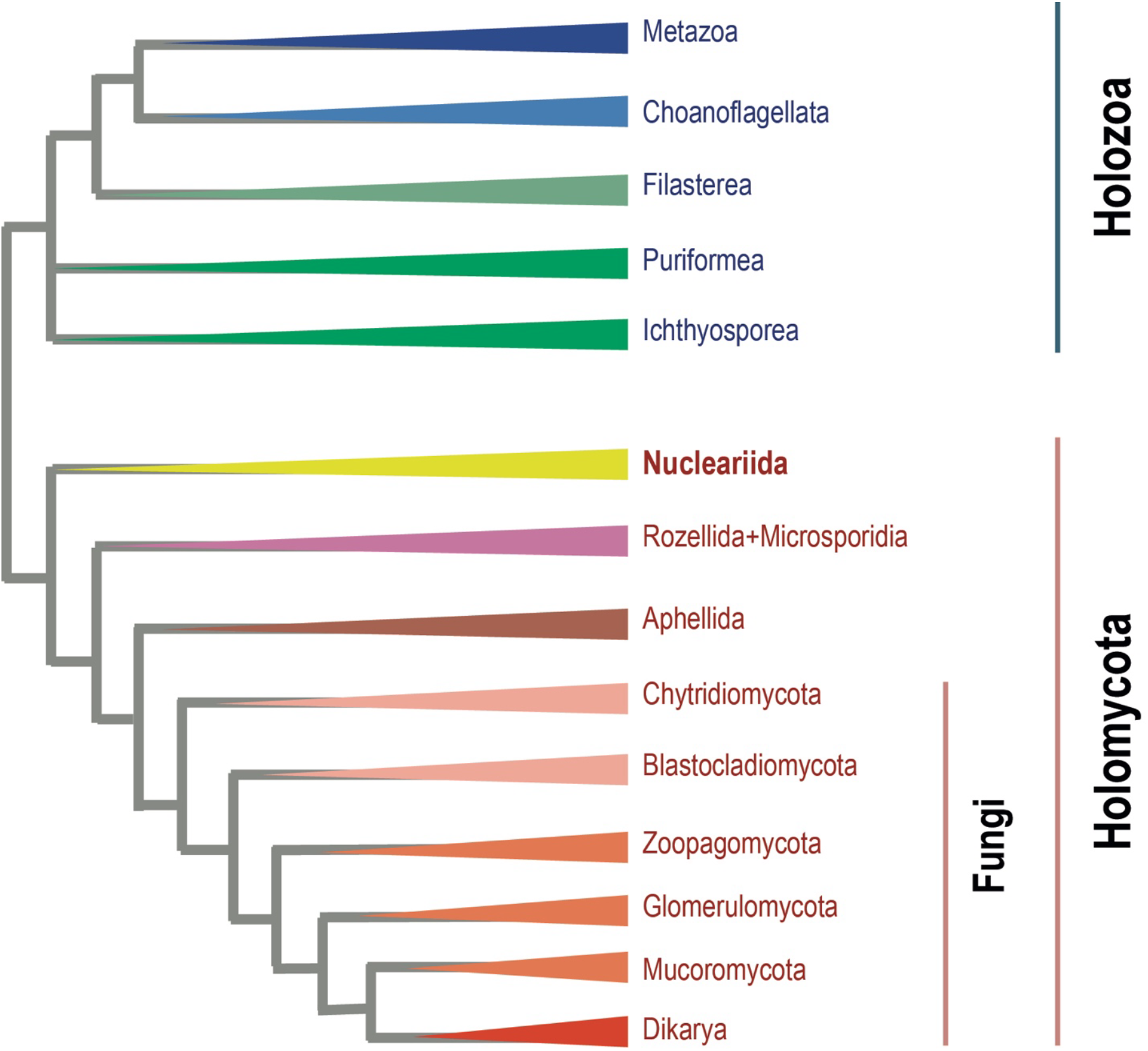
Phylogenetic tree of Opisthokonta. Based on Hehenberger et al., 2017, Galindo et al., 2022, Ocaña-Pallarès et al., 2022

## Materials and methods

### 1. Organism cultivation, DNA extraction, and sequencing

The *N. thermophila* strain …. was isolated from samples of water and sediments from a natural reservoir in the Leningrad district and is kept in culture at the Zoological Institute of the Russian Academy of Sciences (Pozdnyakov et al., 2023). The culture is grown on a monoculture of *E. coli* as food according to the method described in Pozdnyakov et al., 2023.

To reduce the level of bacterial contamination of *N. thermophila* cells, before DNA extraction, the culture liquid was filtered with the washing by PJ medium through 5 μm-pore acetate-cellulose filters on a filtration unit…. The volume of the washing medium exceeded the initial volume of the culture liquid by approximately 25 times. Filters were replaced as they became dirty. *N. thermophylla* cells remained in the liquid above the filter. Filtration was carried out twice, with an interval of 3 days.

Next, the liquid with *N. thermophylla* cells was centrifuged for 1,5 hours at 13300 g in 1.5 μl plastic tubes. *N. thermophila* cells did not settle during centrifugation but were concentrated in the lower layer of liquid.

After centrifugation, the top layer of liquid was removed, and 300 μl of the bottom volume was left in the tube. Next, 300 μl of SDS lysis buffer with a twofold increased concentration of components and…. Proteinase-K were added to it. The finished solution was stirred and kept for three hours at 55 degrees Celsius. Next, the DNA was isolated and purified by phenol-chloroform extraction.

For Illumina HiSeq4000 sequencing, two paired-end libraries were prepared following the TruSeq and Nextera library preparation protocols with an insert length of 700 bp. A total of 900 million paired-end reads were obtained for the two libraries.

The long reads were generated with PromethION sequencing (Oxford Nanopore Technologies, UK). The sequencing libraries were prepared using the ligation sequencing kit SQK-LSK109, native barcoding expansion kit EXP-NBD104 and EXP-NBD114. The gained library for ONT sequencing was then loaded into the flow cells (FLO-MIN106 and FLO-PRO002). Thus, the libraries were obtained with 1 million long reads.

### 2. Genome Assembly and Annotation

The initial genomic assemblies were performed with Flye v.2.9.1 [Kolmogorov et al., 2019] with default settings using long reads library. The draft assembly was checked for contaminations with BlobToolKit v2.3.3 [Challis et al., 2020]. Trusted contigs were selected based on the annotation of contigs against NCBI nucleotide and UniProt reference proteomes databases, GC content, and short/long reads coverage information. Further, the long reads library was mapped on the trusted contigs using the Minimap 2, v.2.24 [Li, 2018], and all unmapped reads were discarded.

The next step assembly was made with the trusted long reads with Flye, v.2.9.1, using the same way. The new assembly was polished in Racon, v.1.4.3 [Vaser et al., 2017] using the selected Illumina paired-end reads. An attempt at the second polishing step in Pilon, v.1.24 [Walker et al., 2014] resulted in a slight deterioration in quality and was rejected. The assembly quality was controlled with Busco, v.5.4.2 [Simão et al., 2015] and QUAST, v.5.0.2 [Gurevich et al., 2013]. on every step of assembly.

Structure annotation of assembled genome was carried out using funannotate pipeline, v.1.8.13 [Palmer, Stajich, 2020], which includes repeat masking with tantan, ab initio gene-prediction training (Augustus, PASA, SNAP, GlimmerHMM, GeneMark), generating consensus gene model (Evidence Modeler) [Haas et al., 2008] and consolidating information from InterProScan v5.65-97.0 and EggNOG annotators, supplemented by additional functional annotating of proteins against several databases (Pfam, InterPro, GO, dbCAN, BUSCO, MEROPS, EggNog, COG). We improved gene prediction with protein evidence from UniProt database. Secreted proteins were estimated with the Phobius command line tool[Käll et al., 2007], and tRNAs were predicted in silico with a tRNAscan-SE algorithm [Chan, Lowe, 2019] included in the funannotate pipeline. We performed structure and functional comparison with genomes of different species of Opisthokonta supergroup (*Fonticula alba* (Holomycota, Rotosphaerida); *Rozella allomycis* (Holomycoa, Rozellida); *Amoeboaphelidium protococcarum* (Holomycota, Aphelida); *Amoeboaphelidium occidentale* (Holomycota, Aphelida); *Gonapodya prolifera* (Holomycota, Chytridiomycota); *Spizellomyces punctatus* (Holomycota, Chytridiomycota); *Blastocladiella emersonii* (Holomycota, Blastocladiomycota); *Allomyces macrogynus* (Holomycota, Blastocladiomycota); *Neurospora crassa* (Holomycota, Dikarya); *Sphaeroforma arctica* (Holozoa, Teretosporea); *Capsaspora owczarzaki* (Holozoa, Filasterea); *Salpingoeca rosetta* (Holozoa, Choanoflagellata); *Monosiga brevicollis* (Holozoa, Choanoflagellata); *Amphimedon queenslandica* (Metazoa, Porifera), *Stylophora pistillata* (Metazoa, Cnidaria) and *Caenorhabditis elegans* (Metazoa, Ecdysosoa)). For that purpose these genomes were augmented with annotations from InterProScan v5.65-97.0 and the funannotate annotate command to ensure a robust basis for comparison. Then the comprehensive analyses were executed using the compare command within the funannotate pipeline. To estimate PFAM motifs occurrence in holomycotan genomes, we used non-metric multidimensional scaling (NMDS) projection of a Bray–-Curtis distance matrix implemented in the funannotate compare.

### 2. Estimation of functional features of genomes based on the presence of PFAM domains

A comparative analysis of the content of domains from the PFAM database was carried out on the basis of the obtained lists of PFAM domains based on annotations.

To arrange species in two-dimensional space based on the functionality of their genomes, we applied a non-metric multidimensional scaling (NMDS) algorithm to functional domain annotations using PFAM domain abundance data (……).

The following sets of domains were selected from the data array of the PFAM domains:

1. Putative unique domains of the common ancestor of Holozoa. It is domains that are absent in all holomycot representatives and present in *S. arctica* and at least in one other representative of Holozoa (since the most basal node is located between Ichthyosporea lineage and other lineages of Holozoa).
2. Putative unique domains of the common ancestor of Holomycota. It is domains that are absent in all holozoan representatives and present in *N. thermophylla* or *F. alba* and at least one other representative of Holomycota (i.e., the most basal node of the Holomycota branch is located between Rotospherida and other lineages).
3. Common domains of *N. thermophila* and Holozoa, lost in other Holomycota. It is domains that are absent in all Holomycota except *N. thermophila* and present in *N. thermophila* and at least one representative of Holozoa.

For each domain from these primary “raw” sets, its taxonomic distribution and cell functions were clarified using the InterPro database. On this ground, the compositions of sets were refined. For putative ancestral domain sets, the newly appeared and the lost domains were identified. A domain was considered lost if itwas absent in one of the Opisthokonta lineages, while present in some other groups of eukaryotes. To provide a more robust comparison we considered a domain to be truly present outside the Opisthokont group if it occurred in at least three lineages and more than two species. If a domain was found in one or two eukaryotic lineages, its presence was considered in the same way.

Finally, all the domains in each of the sets were divided into the following functional categories:

1. Signaling and regulatory functions;
2. Membrane operations and cell adhesion;
3. Protein-protein interactions;
4. Metabolism;
5. Cytoskeleton.
6. Receptor functions;

5) Operations with DNA and the cell cycle;

6) Unknown function.

Then the percentage of domains of each category was calculated for each set and visualized via pie charts.

## Results

The hybrid assembly based on Oxford Nanopore and Illumina paired-end reads yielded an *N. thermophila* genome with a total length of 49 571 131 bp, which were distributed among 571 scaffolds, with an N50 equal to 391 340 and a GC percentage 44.1 (NCBI BioProject accession number ). The assembled genome contains 20.90% of repetitive sequences. The average coverage of Nanopore long reads and Illumina paired-end reads was about 30-fold and 200-fold, respectively. We identified 15 539 genes, of which 15 321 are protein-coding ones. The average lengths of the predicted genes and proteins were 2370.5 nucleotides and 522.3 amino acids, respectively. All genes were functionally annotated by several databases: InterPro (11997), Pfam (10446), GO (10372), EggNOG (6294), BUSCO Eukaryota odb10 (295), MEROPS (396) and dbCAN (229). In addition, we annotated 1731 secreted proteins and 218 tRNA-encoding genes.

The comparison of the obtained assembly with the selected set of genome assemblies of different representatives of Opisthokonta in terms of key indicators showed its comparable characteristics (**Table 1**). On the two-dimensional NMDS diagram, *N. thermophila* is located separately, not grouping with any organisms (**Fig. 2**). At the same time, its closest neighbors (located, however, fairly distantly) are organisms from the Holozoa group.

**Table 1.** Comparison of the genome assemblies of *Nuclearia thermophila* and related species.

|  | Assembly Size | Largest Scaffold | Average Scaffold | Num Scaffolds | Scaffold N50 | Percent GC | Num Genes | Num Proteins | Num tRNA | Unique Proteins | Unique PFAM domains | Prots at least 1 ortholog |
| --- | --- | --- | --- | --- | --- | --- | --- | --- | --- | --- | --- | --- |
| **species** |  |  |  |  |  |  |  |  |  |  |  |  |
| **Nuclearia sp.** | 49,571,131 bp | 968,275 bp | 86,815 bp | 571 | 391,340 bp | 44.10% | 15,539 | 15,321 | 218 | 9,645 | 4,002 | 5,676 |
| **Salpingoeca rosetta** | 55,440,309 bp | 2,682,829 bp | 360,002 bp | 154 | 1,519,553 bp | 52.23% | 11,781 | 11,725 | 115 | 6,729 | 3,924 | 4,994 |
| **Rozella allomycis CSF55** | 11,859,274 bp | 719,121 bp | 11,199 bp | 1,059 | 58,027 bp | 34.48% | 6,350 | 6,350 | 0 | 3,456 | 2,671 | 2,894 |
| **Gonapodya prolifera JEL478** | 48,794,828 bp | 1,572,201 bp | 138,622 bp | 352 | 347,324 bp | 51.75% | 13,911 | 13,831 | 84 | 8,572 | 3,640 | 5,251 |
| **Monosiga brevicollis MX1** | 41,709,928 bp | 3,607,471 bp | 190,456 bp | 219 | 1,073,601 bp | 50.89% | 9,199 | 9,172 | 25 | 4,555 | 3,606 | 4,614 |
| **Neurospora crassa OR74A** | 41,102,378 bp | 9,798,893 bp | 1,957,256 bp | 21 | 6,000,761 bp | 48.19% | 10,544 | 10,784 | 427 | 6,616 | 4,014 | 4,136 |
| **Fonticula alba** | 31,296,464 bp | 4,772,237 bp | 146,245 bp | 214 | 2,529,562 bp | 55.13% | 6,449 | 6,309 | 546 | 3,867 | 2,495 | 2,436 |
| **Amoeboaphelidium protococcarum** | 24,734,778 bp | 3,250,117 bp | 95,871 bp | 258 | 2,170,272 bp | 40.50% | 13,180 | 13,180 | 0 | 6,645 | 3,182 | 6,337 |
| **Spizellomyces punctatus DAOM BR117** | 24,131,112 bp | 2,242,449 bp | 635,029 bp | 38 | 1,465,700 bp | 47.16% | 9,164 | 9,422 | 207 | 3,920 | 4,061 | 5,492 |
| **Capsaspora owczarzaki ATCC 30864** | 27,967,784 bp | 3,794,338 bp | 332,950 bp | 84 | 1,617,775 bp | 52.80% | 8,757 | 8,790 | 106 | 4,273 | 3,953 | 4,510 |
| **Amoeboaphelidium occidentale** | 13,559,732 bp | 366,412 bp | 14,258 bp | 951 | 73,507 bp | 39.93% | 7,568 | 7,495 | 74 | 3,489 | 3,321 | 4,006 |
| **Blastocladiella emersonii ATCC 22665** | 32,654,225 bp | 5,142,960 bp | 1,554,963 bp | 21 | 2,021,495 bp | 65.21% | 10,254 | 10,031 | 223 | 4,399 | 3,406 | 5,515 |
| **Allomyces macrogynus ATCC 38327** | 57,060,573 bp | 3,145,054 bp | 564,956 bp | 101 | 1,114,524 bp | 56.80% | 19,266 | 19,447 | 492 | 10,666 | 3,475 | 8,734 |
| **Amphimedon queenslandica** | 164,261,974 bp | 1,888,931 bp | 12,509 bp | 13,132 | 123,180 bp | 30.98% | 20,057 | 23,125 | 106 | 15,692 | 4,582 | 7,156 |
| **Caenorhabditis elegans** | 100,286,401 bp | 20,924,180 bp | 14,326,629 bp | 7 | 17,493,829 bp | 35.44% | 44,550 | 27,494 | 634 | 21,106 | 4,207 | 5,916 |
| **Nematostella vectensis** | 269,418,438 bp | 22,168,631 bp | 5,732,307 bp | 47 | 17,865,913 bp | 40.71% | 32,275 | 30,801 | 9,571 | 19,524 | 5,395 | 8,449 |
| **Sphaeroforma arctica JP610** | 121,631,465 bp | 357,797 bp | 7,787 bp | 15,619 | 64,598 bp | 33.51% | 18,537 | 18,730 | 318 | 14,929 | 3,310 | 3,796 |

**Figure 2.**
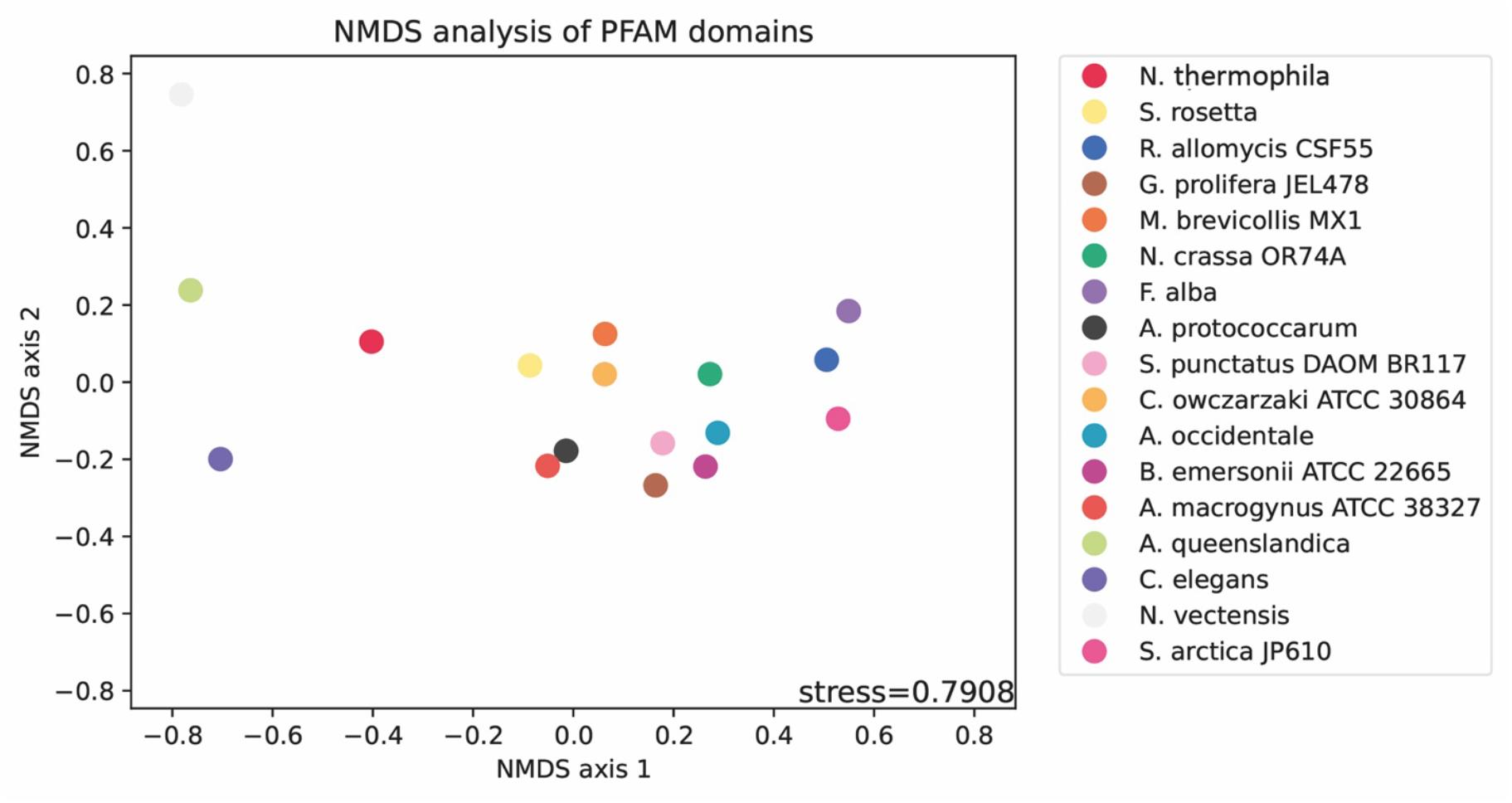
NMDS analysis showing PFAM domain co-occurrence in the genomes of *Nuclearia thermophila* and related species.

The predicted proteome of *N. thermophila* contains 4002 unique PFAM domains, of which 3598 domains are common to representatives of both Holozoa and Holomycota. Out of these, 166 domains are present in members of Holozoa, and *N. thermophila* but absent in other Holomycota (Fig. 3). The distribution of these domains across functional categories shows that the most part (26%) of them are domains associated with signal-regulatory processes. The next most numerous (20%) are domains related to metabolism. The third most numerous (16%) are domains associated with protein-protein interactions. Very noteworthy is the presence of domains associated with cytoskeletal elements characteristic of Holozoa. For instance, PF00435 corresponds to a spectrin domain, PF21033 is involved in the regulation of microtubule dynamics, and PF00038 is associated with the formation of distinctive intermediate filaments.

**Figure 3.**
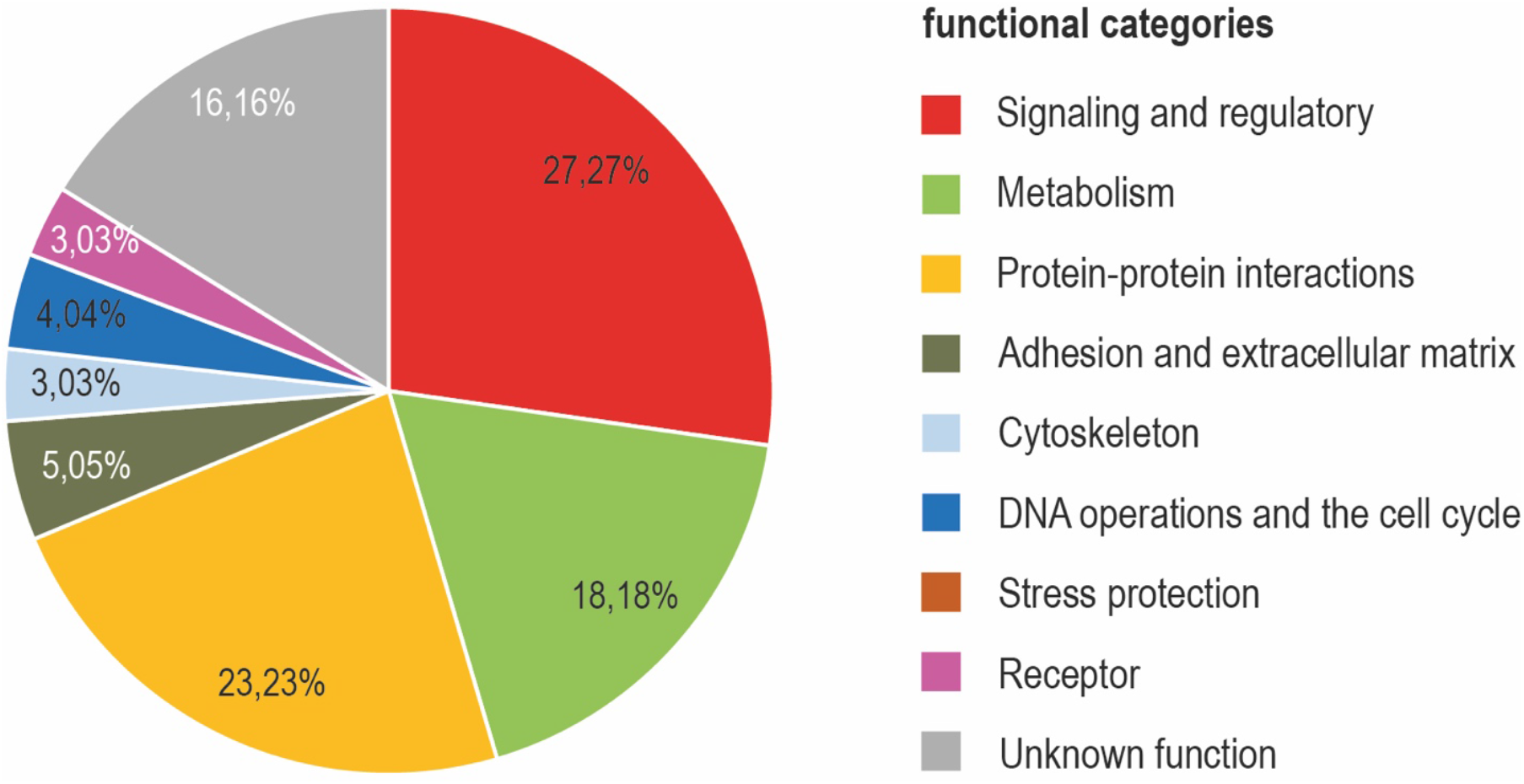
Distribution of PFAM domains common to N. thermophila and Holozoa by functional categories (percentage)

Another 74 domains are shared between *N. thermophila* and other Holomycota and are absent in members of Holozoa (**Supplementary Table 1**). Out of these 74 Holomycota-specific domains 52 are probable Holomycota ampomorphies, and 22 were unique in Opistokontha, because they were lost in Holozoa. Similarly, 85 Holozoa-specific domains were identified, 51 of which were acquired in Holozoa and 34 lost in Holomycota (Fig. 4). Among the domains that appeared in Holomycota, those related to metabolism are predominant. The second largest category is domains of unknown function. The third one are signaling and regulatory domains. Among the likely lost domains in common ancestor of Holomycota, signaling and regulatory domains predominate, and in second place are those related to metabolism and protein interactions.

**Figure 4.**
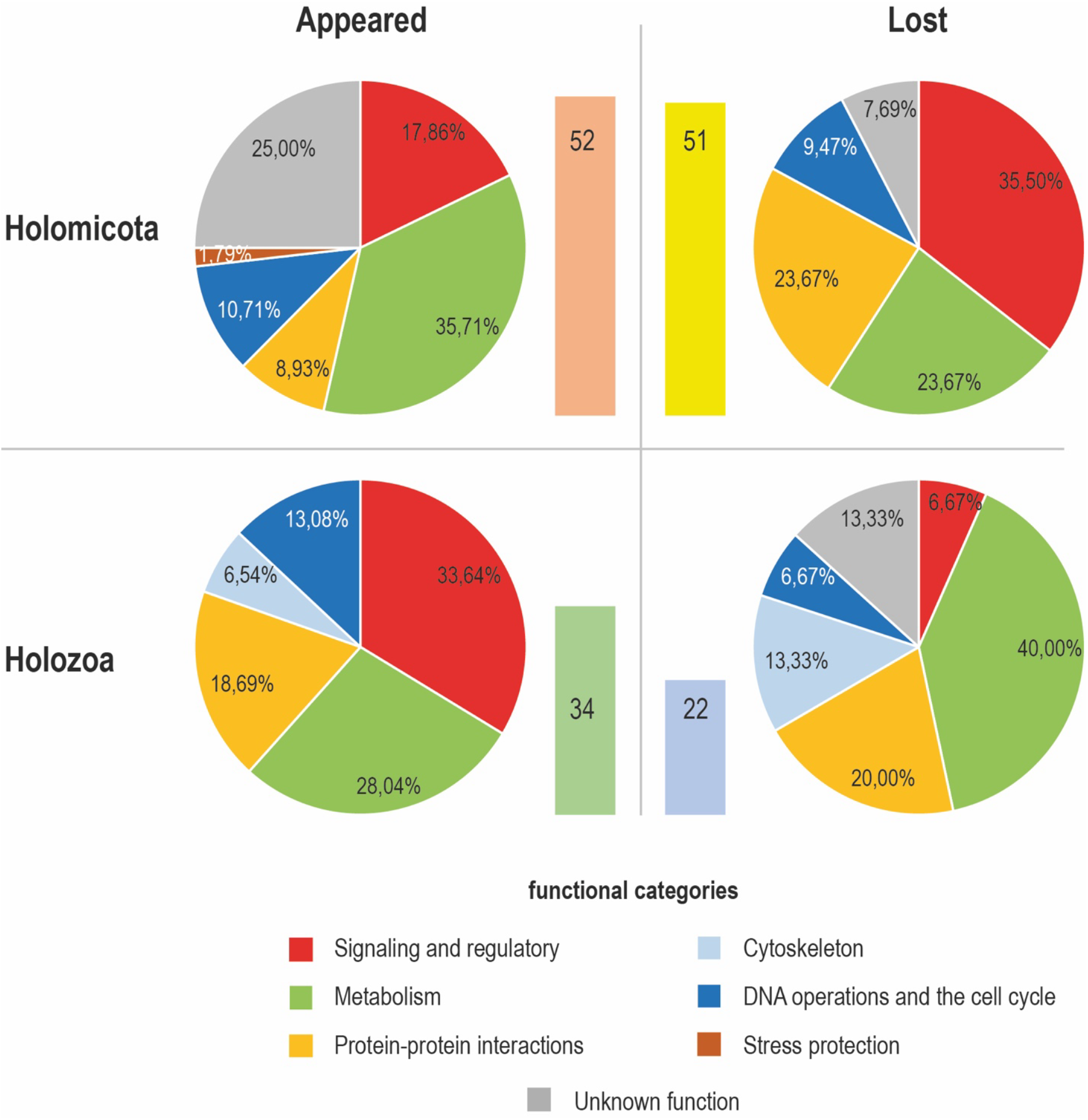
Distribution of PFAM domains presumably acquired or lost in the common ancestors of Holomycota and Holozoa by functional category (percentage). The vertical column is the number of domains.

## Discussion

The genome size of *N. thermophila* is quite large for a haploid unicellular opisthokont, falling within the group of largest opistoknont genomes (40-50 million bp) with the exception of haploid genome of *S. arctica* (122 million bp). Among opistokonts the genome of *N. thermophila* has one of the largest number of protein-coding genes, as well as one of the highest proporsions of the genes that have orthologs in related organisms.

The only previously available sequenced and annotated nucleariid genome is the one of *Fonticula alba*. The number of protein-coding genes and, accordingly, orthogroups found in *N. thermophila* is significantly higher, than in *F. alba* (Table 1). The same is true for PFAM domains shared with other opisthokonts. This may indicate difference between genome assembly strategies applied, but considering specific slime mould life style of *F. alba* this may also be caused by *F. alba* reductive evolution. Being more derived and secondarily modified representative of nucleariids *F. alba* could undervent a considerable gene loss.

According to the integrated indicator of the number and diversity of PFAM domains, N. thermophila shows similarity with free-living representatives of Holozoa (Fig. 2). This correlates with the large number of discovered unique PFAM domains. It is worth noting here that F. alba is close to R. allomycis and S. arctica, and all three of them are the most distant from N. thermophila and its neighbors. Obviously, the indicators of the number and diversity of PFAM domains for them are minimal.

After subtracting the common opisthokont domains, N. thermophila still has twice as many PFAM domains common with Holozoa as domains common with the rest of Holomycota. Therefore, despite belonging to Holomycota clade, nucleariids still share a lot of genomic traints with Holozoa, which most probably can be interpreted as inherited from the common opistokonta ancestor. These results indicate that during the early stages of Holomycota evolution their metabolic repertoire, signaling mechanisms and protein interactions remained relatively conserved and were drastically modified only in the course of evolution of more derived holomycotans.

It is likely that the common ancestor of opistoknota was capable of both amoeboid and flagellar locomotion, possible swithching between the two stages in the course of life cycle or environmental condition. This feature is still retained in some free living Holozoa (e. g. Syssomonas spp. and Pygoraptor spp.) (Hehenberger et al., 2017). Partly similarity of nucleariids to Holozoa in signaling, protein interactions and special cytoskeletal elements can be explain by reaining the same ancestral amoeboid “toolkit”, which was also inherited by Holozoa.

The obtaining Obtaining a new nucleariid genome also allows comparison of gains and losses of protein domains between Holozoa and Holomycota ancestors (Fig. 4). This analysis suggested that, perhaps compared with the ancestor of Holozoa, the ancestor of Holomycota underwent much more dramatic deviations from the hypothetical common ancestor of Opistokontha. The number of both gains and losses of protein domains is approximately twice as high: the ancestor Holomycota has 52 acquired domains and 51 lost; The ancestor Holozoa acquired 34 domains and lost 22. In addition, from our analysis the idea can be appeared that in the ancestor of Holomycota, most of the gained domains related to metabolism, and in the ancestor of Holozoa it related to signaling and regulation. If so, it is consistent with recently identified trends in the evolution of the Holozoa and Holomycota groups (Ocaña-Pallarès et al., 2022) However, conclusions from the noted observations can still only be assumptions. In our analysis, only those domains present in S. arctica were noted as ancestral domains of Holozoa, but S. arctica shows a reduced diversity of PFAM domains compared to the rest Holozoa (**Table 1, Fig. 2**), probably due to its prasitic lifestyle. To confirm the noted trends, it is necessary to study the genome of a free-living representative of the group Teretosporea, for example, Sissomonas sp.

## Acknowledgements

The authors are grateful to Sergey A. Karpov, Alexey V. Smirnov and Elena S. Nassonova for advice on various research-related issues, the team of Invertebrate Zoologe Department of SPbSU for colaboration and the staff of the Zoological Institute of the Russian Academy of Sciences for creating comfortable working conditions.

## References

1. Adl SM, Bass D, Lane CE, Lukeš J, Schoch CL, Smirnov A, Agatha S, Berney C, Brown MW, Burki F, Cárdenas P, Cěpička I, Chistyakova L, del Campo J, Dunthorn M, Edvardsen B, Eglit Y, Guillou L, Hampl V, Heiss AA, Hoppenrath M, James TY, Karnkowska A, Karpov S, Kim E, Kolisko M, Kudryavtsev A, Lahr DJG, Lara E, Le Gall L, Lynn DH, Mann DG, Massana R, Mitchell EAD, Morrow C, Park JS, Pawlowski JW, Powell MJ, Richter DJ, Rueckert S, Shadwick L, Shimano S, Spiegel FW, Torruella G, Youssef N, Zlatogursky V, Zhang Q. Revisions to the classification, nomenclature, and diversity of eukaryotes. J Eukaryot Microbiol. 2019, 66:4–119.

2. Amaral-Zettler L.A,. Nerad T.A., O’Kelly C.J., Sogin M.L. The nucleariid amoebae: more protists at the animal-fungal boundary. J Eukaryot Microbiol. 2001, 48:293–297.

3. Brown MW, Spiegel FW, Silberman JD (2009) Phylogeny of the “forgotten” cellular slime mold, Fonticula alba, reveals a key evolutionary branch within Opisthokonta. Mol Biol Evol 26:2699–2709.

4. Camacho C., Coulouris G., Avagyan V., Ma N., Papadopoulos J., Bealer K., Madden TL. BLAST+: Architecture and applications. BMC Bioinform. 2009, 10, 421. 10.1186/1471-2105-10-421.

5. Challis, R.; Richards, E.; Rajan, J.; Cochrane, G.; Blaxter, M. BlobToolKit–interactive quality assessment of genome assemblies. G3-Genes Genom. Genet. 2020, 10, 1361–1374. 10.1534/g3.119.400908.

6. Chan, P.P.; Lowe, T.M. tRNAscan-SE: Searching for tRNA Genes in Genomic Sequences. In Gene Prediction. Methods in Molecular Biology; Kollmar, M., Ed.; Humana: New York, NY, USA, 2019; Volume 1962. 10.1007/978-1-4939-9173-0_1.

7. Cienkowski L (1865) BeiträgezurKenntnisderMonaden. ArchmikroskAnat1:203–232.

8. Gabaldón T., Völcker E., Torruella G. On the Biology, Diversity and Evolution of Nucleariid Amoebae (Amorphea, Obazoa, Opisthokonta). Protist. 2022, 173(4):125895. Doi: 10.1016/j.protis.2022.

9. Galindo LJ., Torruella G., López-García P., Ciobanu M., Gutiérrez-Preciado A., Karpov SA., Moreira D. Phylogenomics supports the monophyly of aphelids and fungi and identifies new molecular synapomorphies. Syst. Biol. 2022, 72:505–515. Doi: 10.1093/sysbio/syac054.

10. Glockling S.L., Marshall W.L., Gleason F.H. Phylogenetic interpretations and ecological potentials of the Mesomycetozoea (Ichthyosporea). Fungal Ecology. 2013, 6(4), 237-247. Doi 10.1016/j.funeco.2013.03.005.

11. Gurevich, A.; Saveliev, V.; Vyahhi, N.; Tesler, G. QUAST: Quality assessment tool for genome assemblies. Bioinformatics 2013, 29, 1072–1075. 10.1093/bioinformatics/btt086.

12. Haas, B.J.; Salzberg, S.L.; Zhu, W.; Pertea, M.; Allen, J.E.; Orvis, J.; White, O.; Buell, C.R.; Wortman, J.R. Automated eukaryotic gene structure annotation using EVidenceModeler and the Program to Assemble Spliced Alignments. Genome Biol. 2008, 9, R7. 10.1186/gb-2008-9-1-r7.

13. Hehenberger, Elisabeth; Tikhonenkov, Denis V.; Kolisko, Martin; Campo, Javier del; Esaulov, Anton S.; Mylnikov, Alexander P.; Keeling, Patrick J. (2017). “Novel Predators Reshape Holozoan Phylogeny and Reveal the Presence of a Two-Component Signaling System in the Ancestor of Animals”. Current Biology. 27 (1): 2043–2050.e6. doi:10.1016/j.cub.2017.06.006. PMID 28648822.

14. Hertel L. A.; Bayne C. J.; Loker, E. S. (August 2002), “The symbiont Capsaspora owczarzaki, nov. gen. nov. sp., isolated from three strains of the pulmonate snail Biomphalaria glabrata is related to members of the Mesomycetozoea”, International Journal for Parasitology, 32 (9): 1183–91, doi:10.1016/S0020-7519(02)00066-8, PMID 12117501.

15. Hess S., Suthaus A. The vampyrellid amoebae (Vampyrellida, Rhizaria). Protist. 2022, 173:125854. Doi: 10.1016/j.protis.2021.125854.

16. Jones, P.; Binns, D.; Chang, H.-Y.; Fraser, M.; Li, W.; McAnulla, C.; McWilliam, H.; Maslen, J.; Mitchell, A.; Nuka, G.; et al. InterProScan 5: Genome-scale protein function classification. Bioinformatics 2014, 30, 1236–1240. 10.1093/bioinformatics/btu031.

17. Käll, L.; Krogh, A.; Sonnhammer, E.L. Advantages of combined transmembrane topology and signal peptide prediction—The Phobius web server. Nucleic Acids Res. 2007, 35, W429–W432. 10.1093/nar/gkm256.

18. Kolmogorov, M.; Yuan, J.; Lin, Y.; Pevzner, P. Assembly of Long Error-Prone Reads Using Repeat Graphs. Nat. Biotechnol. 2019, 37, 540–546. 10.1038/s41587-019-0072-8.

19. Li, H. Minimap2: Pairwise alignment for nucleotide sequences. Bioinformatics 2018, 34, 3094–3100. 10.1093/bioinformatics/bty191.

20. Liu Y, Steenkamp ET, Brinkmann H, Forget L, Philippe H, Lang BF (2009) Phylogenomic analyses predict sistergroup relationship of nucleariids and fungi and paraphyly of zygomycetes with significant support. BMC Evol Biol 9: 1–11.

21. Medina M., Collins A.G., Taylor J.W., Valentine J.W., Lipps J.H., Amaral-Zettler L., Sogin M.L. Phylogeny of Opisthokonta and the evolution of multicellularity and complexity in Fungi and Metazoa. Int J Astrobiol. 2003, 2:203–211

22. Ocaña-Pallarès E., Williams TA., López-Escardó D. et al. Divergent genomic trajectories predate the origin of animals and fungi. Nature 2022, 609:747–753. Doi: 10.1038/s41586-022-05110-4

23. Palmer, J.; Stajich, J. Nextgenusfs/Funannotate: Funannotate v1.8.1, Version 1.8.1; Zenodo: 2020. 10.5281/zenodo.4054262.

24. Patterson, D.J., Nygaard, K., Steinberg, G. & Turley, C.M. (1993). Heterotrophic flagellates and other protists associated with oceanic detritus throughout the water column in the mid North Atlantic. Journal of the Marine Biological Association of the United Kingdom 73: 67–95.

25. Ros-Rocher N., Pérez-posada A., Leger M.M., Ruiz-Trillo I. The origin of animals: an ancestral reconstruction of the unicellular-to-multicellular transition. Open Biol. 2021, 11:200359.

26. Simão, F.A.; Waterhouse, R.M.; Ioannidis, P.; Kriventseva, E.V.; Zdobnov, E.M. BUSCO: Assessing genome assembly and annotation completeness with single-copy orthologs. Bioinformatics 2015, 31, 3210–3212. 10.1093/bioinformatics/btv351.

27. Tedersoo L., Sánchez-Ramírez S., Kõljalg U., Bahram M., Döring M., Schigel D., May T., Ryberg M., Abarenkov K. High-level classification of the Fungi and a tool for evolutionary ecological analyses. Fungal Diversity. 2018, 90 1):135–159. doi:10.1007/s13225-018-0401-0.

28. Torruella G., de Mendoza A., Grau-Bove X., Anto M., Chaplin MA., del Campo J., Eme L., Perez-Cordo G. Whipps CM., Nichols KM., Paley R., Roger AJ., Sitj-Bobadilla A., Donachie S., Ruiz-Trillo I. Phylogenomics reveals convergent evolution of lifestyles in close relatives of animals and fungi. Current Biology. 2015, 25:2404–2410. DOI: 10.1016/j.cub.2015.07.053

29. Vaser, R.; Sović, I.; Nagarajan, N.; Šikić, M. Fast and accurate de novo genome assembly from long uncorrected reads. Genome Res. 2017, 27, 737–746. 10.1101/gr.214270.116.

30. Walker, B.J.; Abeel, T.; Shea, T.; Priest, M.; Abouelliel, A.; Sakthikumar, S.; Cuomo, C.A.; Zeng, Q.; Wortman, J.; Young, S.K.; et al. Pilon: An Integrated Tool for Comprehensive Microbial Variant Detection and Genome Assembly Improvement. PLoS ONE 2014, 9, e112963. 10.1371/journal.pone.0112963.

31. Westheide, Rieger. 2011

